## Supplemental material containing Figs. S1-S3 for "The MAPK/ERK channel capacity exceeds 6 bit/hour"

**CONTENTS**

**S1 Fig.** Experimental and data analysis workflow (reconstruction-based approach).

**S2 Fig.** Interval encoding: input sequence reconstruction algorithm and determination of the sources of information loss (reconstruction-based approach).

**S3 Fig.** Cell-to-cell bitrate variability and exclusion of non-responding cells (reconstruction-based approach).

Other supporting materials for this article include:

**Source data** at DOI: 10.5281/zenodo.7808385
**Source code** at URL: https://github.com/pawelnalecz/pulsatile-information

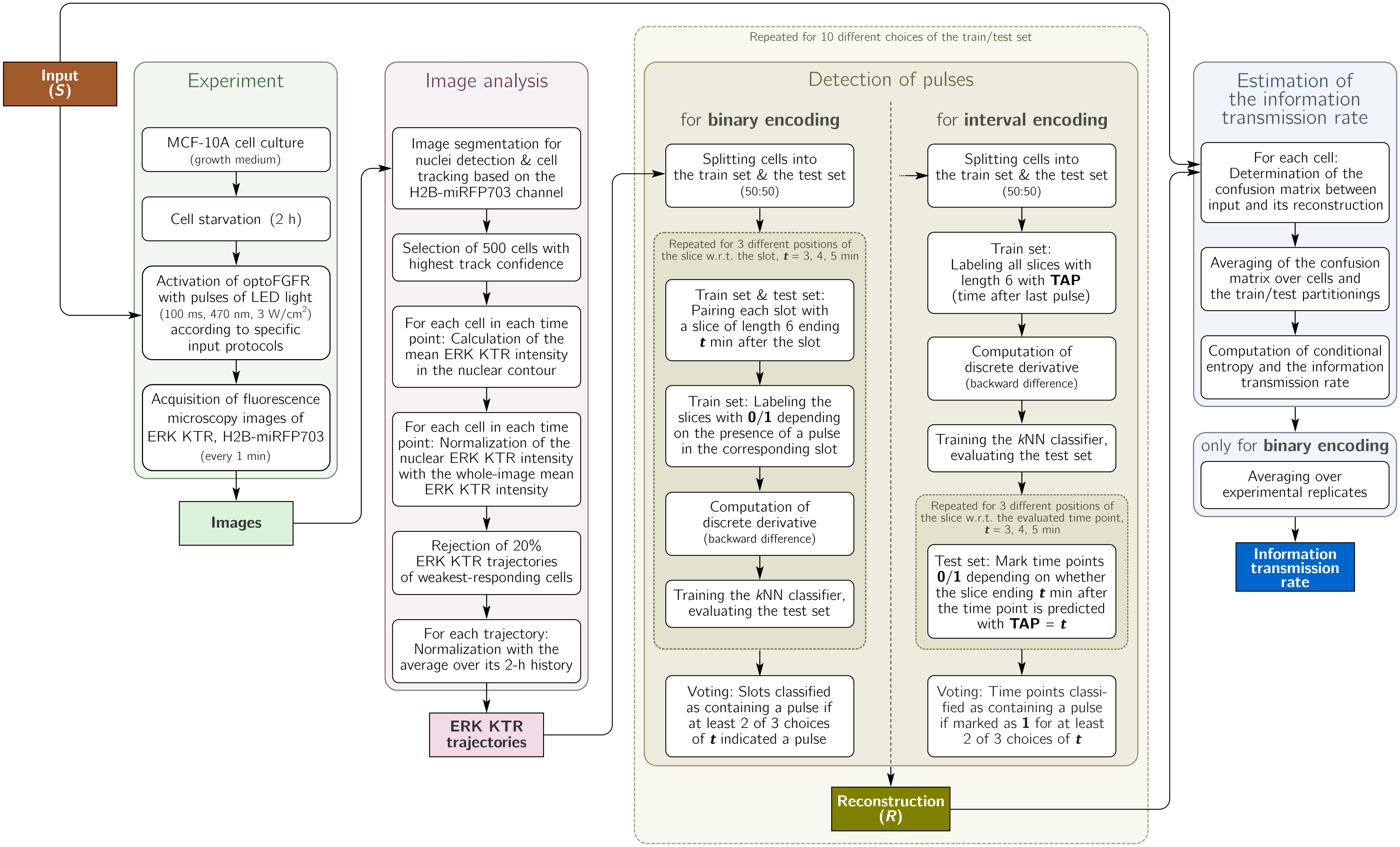


**S1 Fig. Experimental and data analysis workflow (reconstruction-based approach).**

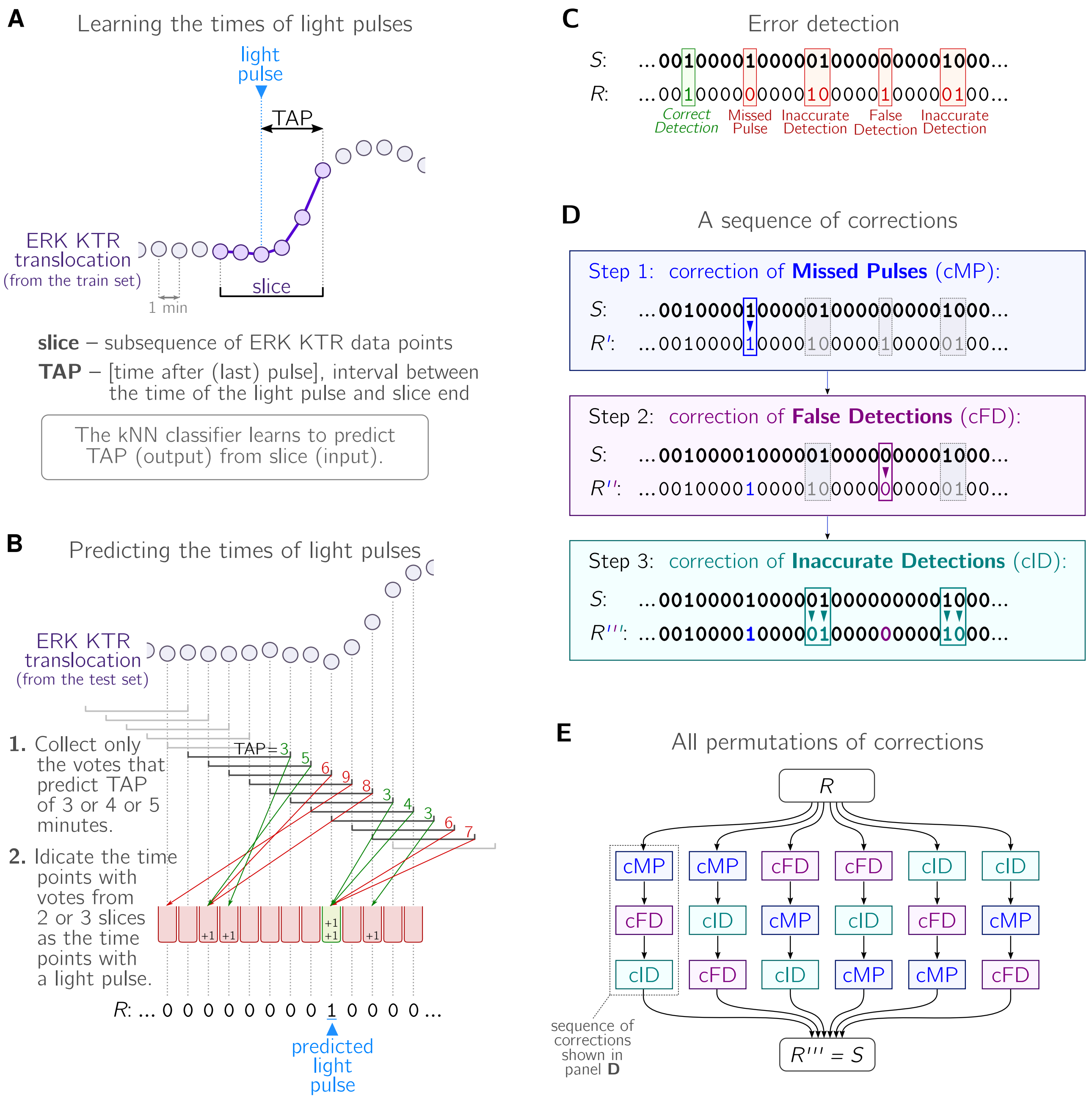


**S2 Fig. Interval encoding: input sequence reconstruction algorithm and determination of the sources of information loss (reconstruction-based approach).**

(**A**) Tracks from the training set are segmented into overlapping slices of 6 consecutive time points such that each time point belongs to 6 slices. Each slice is labeled with the time after pulse (TAP), measured with respect to the last time point in the slice. The *k*NN classifier is trained to predict the TAP based on 5 discrete differences between 6 consecutive time points of a slice.

(**B**) For each slice, the TAP predicted by the classifier indicates a time point at which the stimulation pulse could occur. Votes for particular time points from different slices are counted. Only votes from slices predicted with TAP = 3,4,5 min are taken into account, the remaining are ignored as unreliable. Time points that received at least two out of the three possible votes are considered as time points with pulse in the final reconstruction, *R*.

(**C)** Detection and labeling of errors in the reconstruction *R*. Inaccurate detections (one minute before or after the pulse) are not decomposed into missed pulses and false detections but assigned to their specific error types. Since patterns of ‘11’ and ‘101’ are guaranteed never to occur in the input sequence *S*, the error classification is unambiguous.

(**D**) An example 3-step sequence of error corrections for the interval encoding protocols. After the third step, the fully corrected reconstruction *R*’’’ is identical to the input sequence *S*. The difference between bitrate before and after each correction step is attributed to the particular information loss source.

(**E**) As the difference in bitrate depends on the sequence of corrections, the contributions of the three types of errors are averaged over all permutations of correction steps.

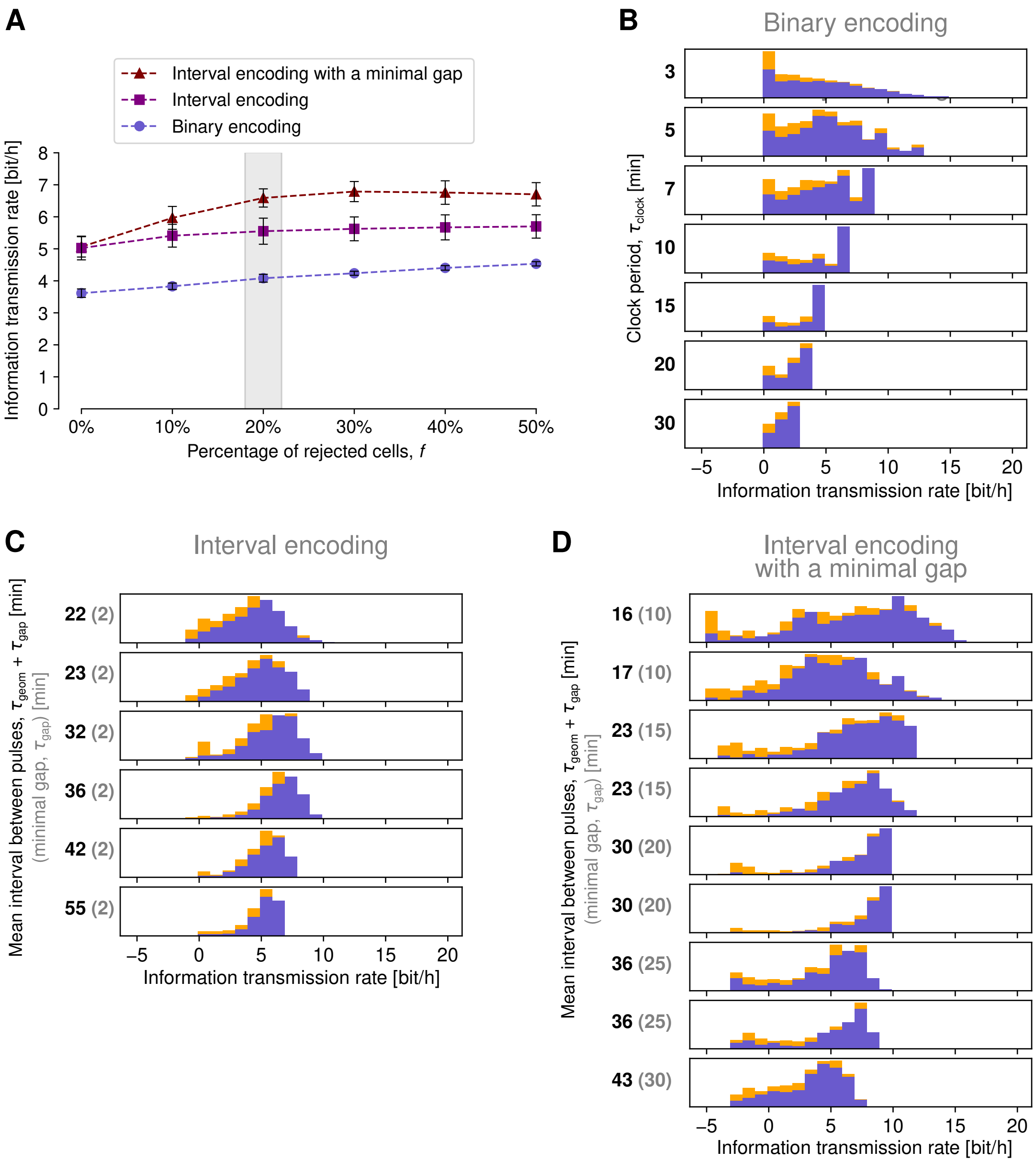


**S3 Fig. Cell-to-cell bitrate variability and exclusion of non-responding cells (reconstruction-based approach)**.

(**A**) Bitrate estimate as a function of the fraction of cells excluded in the preselection step. The (lower bound for) bitrate is computed as in Figure 3. Error bars denote standard error of the mean based on 2–4 experiments with the highest bitrate. In the preselection step we aim at excluding cells that do not respond to stimulation, possibly due to low expression of optoFGFR or ERK KTR. The rejection criterion is formulated in a way *a priori* independent of the accuracy of pulse detection, see Methods for details. Throughout the paper, the fraction of rejected cells is set to 20% (highlighted in gray), because above this value the bitrate estimates in the interval encoding protocols (with and without minimal gap) reach a plateau.

(**B**, **C**, **D**) Histograms of the information transmission rates in single cells for (B) binary encoding, (C) interval encoding, and (D) interval encoding with a minimal gap. Estimates for cells rejected in the preselection step (20% of all cells) are marked in orange. Negative bitrate estimates can occur due to the rough approximation based on inequalities in Eq. (8) in the main text.
